# Dissolved organic matter fluxes and fatty acid composition from shallow-water and deep-water demo- and hexactinellid sponges from New Zealand

**DOI:** 10.1101/2024.06.03.597081

**Authors:** Tanja Stratmann, Mario L. M. Miranda, Anna de Kluijver, Kathrin Busch, Michelle Kelly, Sadie Mills, Peter J. Schupp

## Abstract

Sponges are an important component of shallow- and deep-water ecosystems enhancing eukaryotic biodiversity via diverse endo- and epibiota and by providing three dimensional habitats for benthic invertebrates and fishes. Sponge biodiversity is particularly high in the waters around New Zealand (Southwest Pacific), where we collected two shallow- and two deep-water sponge species (*Tedania* sp., *Suberea meandrina*, *Farrea raoulensis*, *Artemisina* sp.) for *ex-situ* incubation experiments to detect whether these sponges may process dissolved organic matter.

We assessed the biochemical and phospholipid-derived fatty acids (PLFAs) and measured dissolved organic carbon (DOC) and total dissolved nitrogen (TDN) fluxes. Changes in fluorescent dissolved organic matter (FDOM) over time were analyzed and the results were linked to the bacterial communities of the sponge holobiont.

Dried sponge tissue consisted of 17.5 ± 3.75% organic (org.) C and of 4.34 ± 1.02% total nitrogen (TN) with a natural stable isotope composition of -19.0 ±0.25‰ for δ^13^org. C and of 10.2 ±2.43‰ for δ^15^TN. None of the DOC fluxes was significant and only the release of TDN by *Tedania* sp. was significantly different from 0 μmol TDN g org. C^-1^ d^-1^. We detected the presence of four fluorophores in the FDOM pool: 2 tryptophan- and protein-like fluorophores (C1, C2), 1 humic-like fluorophore (C3), and 1 tyrosine-like fluorophore (C4). The maximum fluorescence intensity F_max_ of C1 decreased significantly in *S. meandrina* incubations, whereas F_max_ of C2 grew in the same incubations. F_max_ of C3 increased in *Tedania* sp. incubations, in which F_max_ of C4 decreased. In comparison, F_max_ of C4 in *S. meandrina* rose.

The PLFA composition of sponge tissue was dominated by long-chain fatty acids, saturated fatty acids, and monosaturated fatty acids and most PLFAs were sponge- and bacteria-specific. The bacterial community of the demosponges *Artemisina* sp., *S. meandrina* and *Tedania* sp. consisted mostly of Proteobacteria and Chloroflexota, whereas the dominating bacteria phylum of the hexactinellid *F. raoulensis* was Proteobacteria.

We proposed that the holobionts of *S. meandrina* and *Tedania* sp. contain bacteria that are involved in the transformation and degradation of DOM. In *S. meandrina*, Chloroflexota and Poribacteria may degrade tryptophan-like fluorophores to a chemically modified tryptophan-like and protein-like fluorophore, while producing a tyrosine-like fluorophore. In *Tedania* sp., Chloroflexota may contribute to the release of significant amounts of TDN by producing humic-like fluorophores, while degrading tyrosine-like fluorophores. *Farrea raoulensis* may not take up DOM due to a lack of Poribacteria and Chloroflexota or may use colloidal instead of truly dissolved DOC.

## 1. Introduction

Marine sponges (phylum Porifera) are mostly suspension-feeding, sessile metazoans (though see (Hestetun et al., 2016; Morganti et al., 2021; Vacelet and Boury-Esnault, 1995) whose global distribution stretches from the Arctic (e.g., (Morganti et al., 2022; Rybakova et al., 2020; Stratmann et al., 2022)) to the Southern Ocean (e.g., (Downey et al., 2012; Kersken et al., 2014)). They occur in all water depths from the intertidal zone (e.g., (Barnes, 1999; Gastaldi et al., 2018)) to deep-sea abyssal plains (e.g., (Beaulieu, 2001; Kersken et al., 2019)), and may contribute substantially to benthic density and biomass in sponge grounds (Cathalot et al., 2015; Hawkes et al., 2019; Maldonado et al., 2017; McIntyre et al., 2016; Morganti et al., 2022; Ramiro-Sánchez et al., 2019) and in tropical coral reefs (De Goeij et al., 2017; Diaz and Rützler, 2001). Globally, the highest sponge biodiversities can be found in the northern Gulf of Mexico, the temperate Northern Atlantic including the Mediterranean Sea, the Central Indo-Pacific, and temperate Australasia (Spalding et al., 2007; Van Soest et al., 2012). Waters around New Zealand, in particular, host a minimum of 1361 different species (Kelly and Sim-Smith, 2023) and several new species have been described in recent years (e.g., (Dohrmann et al., 2023; Reiswig et al., 2021)).

Suspension-feeding sponges filter bacterioplankton (de Goeij et al., 2008b; Scheffers et al., 2004; Yahel et al., 2006) and particulate organic carbon (POC) (Hadas et al., 2009; McMurray et al., 2016) out of the water column with pumping rates (ml s^-1^ ml^-1^ sponge) of 0.03 (*Cinachyrella cavernosa* (Lamarck, 1815)) to 17.3 (*Aphrocallistes vastus* Schulze, 1886; (Dahihande and Thakur, 2019; Leys et al., 2011)) and filtration efficiencies of 23% (*Rhopaloeides odorabile* Thompson, Murphy, Bergquist & Evans, 1987) to 99% (*Sarcotragus spinosulus* Schmidt, 1862) for bacteria (Massaro et al., 2012; Trani et al., 2021). Several sponge species, such as *Cliona orientalis* Thiele, 1900, *Ircinia felix* (Duchassaing & Michelotti, 1864), or *Vazella pourtalesii* (Schmidt, 1870), also consume dissolved organic matter (DOM) (Achlatis et al., 2019; Archer et al., 2017; Bart et al., 2020) with DOM removal rates (μmol C g DW_sponge_^-1^ h^-1^) of 3.70±0.26 (*Geodia barretti* Bowerbank, 1858) to 56.1±19.9 (*Acantheurypon spinispinosum* (Topsent, 1904)) (Bart et al., 2020).

DOM is operationally defined as organic matter that passes through filters with pore sizes of 0.2 to 0.7 μm and includes fatty acids, vitamins, sugars, amino acids, proteins, lignins, and polysaccharides (Repeta, 2015). About 90% of the DOM pool, however, is undescribed and consists of complex degradation products of unknown origin (Repeta, 2015). The carbon-containing fraction of DOM is called dissolved organic carbon (DOC) and the nitrogen-containing fraction dissolved organic nitrogen (DON). DOC is the largest organic C pool in the oceans (662 Pg C; (Hansell et al., 2009)) and consists of labile, semi-labile and semi-refractory, and refractory DOC. Labile DOC has a turnover of minutes to days (Fuhrmann and Ferguson, 1986; Keil and Kirchman, 1999) and contributes only <0.2 Pg C (= 0.03% of global DOC pool) (Repeta, 2015). Semi-labile and semi-refractory DOC contribute 20 Pg C (= 3% of global DOC pool) (Hansell et al., 2012) with a turnover of months to years (Hansell et al., 2012) and refractory DOC (including refractory and ultra-refractory DOC) is the largest pool with >642 Pg C (= 97% of global DOC pool) (Hansell et al., 2012) and a turnover of centuries to millennia (Bauer et al., 1992; Druffel et al., 2019). Parts of DOM is colored, the so-called colored dissolved organic matter (CDOM), and absorbs light of the visible and UV spectrum (Coble, 2007). A fraction of this CDOM in turn exhibits blue fluorescence, the so-called fluorescent DOM (FDOM), and may serve as an “optical marker” (Stedmon and Nelson, 2015) for protein-like and humic-like fluorescing compounds (Coble, 2007).

Fatty acids, building blocks of lipids, are more diverse in sponges than in other aquatic fauna (Rod’kina, 2005). Their specific composition varies among sponge classes and is more similar in Hexactinellida and Demospongiae than in Calcarea (Thiel et al., 2002a). Hexactinellida, like *Acanthascus* (*Staurocalyptus*) sp. Ijima, 1897 or *Acanthascus* sp. Schulze, 1886, contain high concentrations of C30Δ^5,9,23^, whereas Demospongiae (e.g., *Spirastrella* sp. Schmidt, 1868, *Astrosclera willeyana* Lister, 1900, *Phakellia ventilabrum* (Linnaeus, 1767)) have high concentrations of C26:1Δ^5,9^(Thiel et al., 2002a). In comparison, unsaturated long-chain fatty acids (LCFA, ≥C24) are absent in Calcarea sponges, such as *Sycon* sp. Risso, 1827 or *Leucetta* sp. Haeckel, 1872 (Thiel et al., 2002b). Sponges produce these LCFA via elongation of fatty acids with shorter chain length that originate from their diets (Carballeira et al., 1986; de Kluijver et al., 2021). Iso (*i*) and anteiso (*ai*)-branched fatty acids and mid-chain branched fatty acids (MBFAs), that are also found in sponges, however, are produced by their bacterial symbionts (Kaneda, 1991; Thiel et al., 1999).

As holobionts (Webster and Taylor, 2012), up to 38% of the sponge volume can consist of microbial cells (Vacelet, 1975). Sponge species with particularly high densities of symbiotic microorganisms (10^8^ to 10^10^ microorganisms g^-1^ sponge tissue; (Hentschel et al., 2006)) are known as high-microbial abundance (HMA) sponges, whereas sponges with symbiotic microbial densities comparable to densities of microorganisms in seawater are known as low-microbial abundance sponges (LMA) sponges (10^5^ to 10^6^ bacteria g^-1^ sponge tissue; (Hentschel et al., 2006)). LMA sponges often host a bacterial core community dominated by a single bacteria clade that differs among sponge specimens (Erwin et al., 2015; Giles et al., 2013). In comparison, HMA sponges host a bacterial community that is more diverse and clearly different from the bacterial community of LMA sponges (Erwin et al., 2015; Giles et al., 2013).

In this study, we measured *ex-situ* DOM fluxes in shallow-water (6 m water depth) and deep-sea sponges (828 to 1448 m water depth) around New Zealand in the Southeast Pacific. We investigated whether DOC and total dissolved nitrogen (TDN) fluxes differed between sponge clusters, and we tried to decipher FDOM components present in the incubations. Furthermore, we studied differences in fatty acid and bacterial composition among the various sponge species.

## 2. Materials and methods

### 2.1 Sampling

Shallow-water sponges were collected by snorkeling at 6 m water depth at Denham Bay (Raoul Island, New Zealand; Fig. 1, Table 1). All sponges were transported in plastic bags filled with ambient seawater to R/V *Sonne* that stayed outside the 1 km exclusion zone around the Kermadec Islands. Deep-water sponges were collected at water depths between 828 m and 1448 m (Fig. 1, Table 1) with the remotely operated vehicle ROV Kiel 6000 (Geomar, Kiel, Germany) during RV *Sonne* cruise SO254 (Simon, 2017) and transported in bioboxes filled with ambient seawater to R/V *Sonne*.

**Figure 1.**
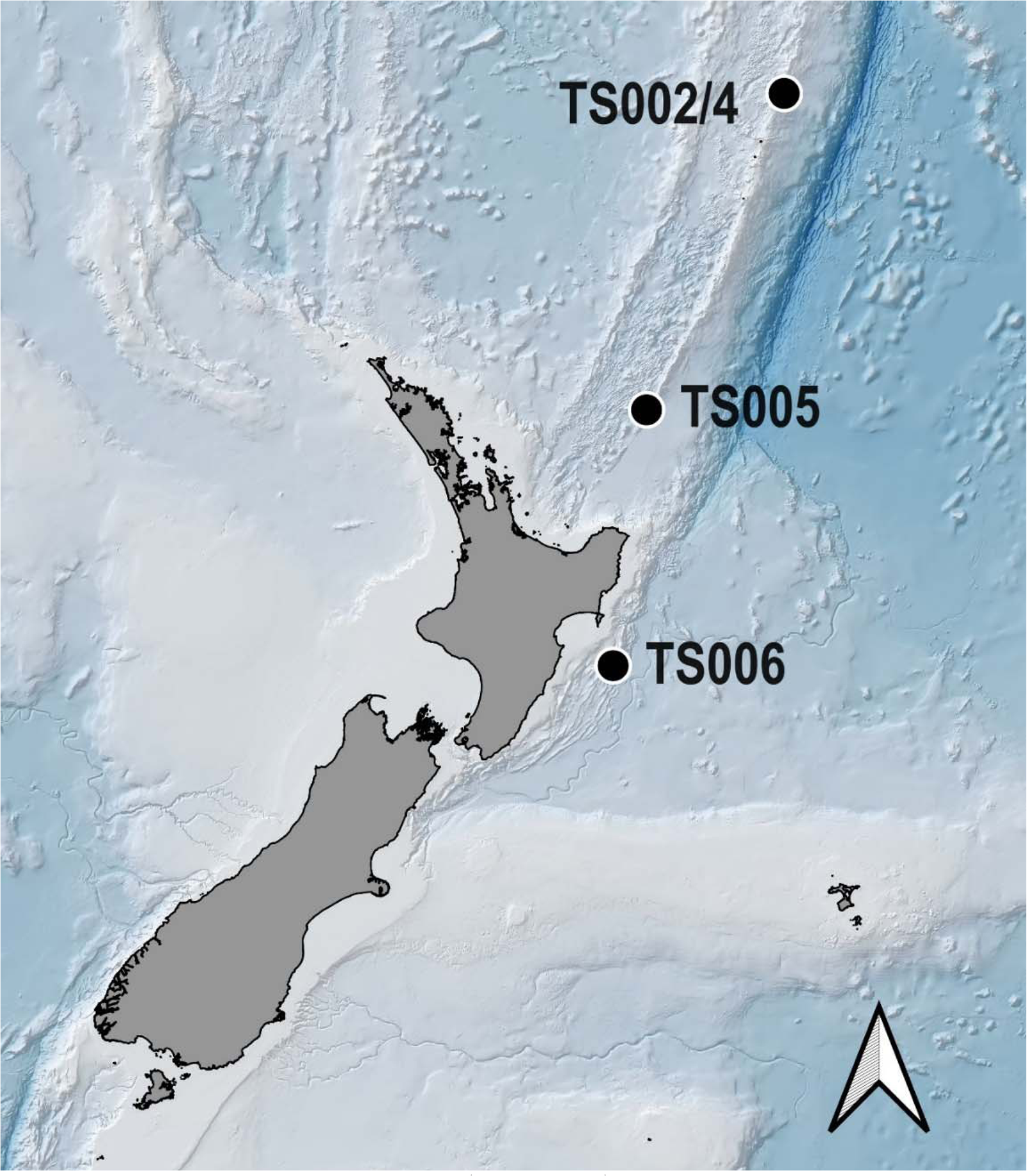
Map with sampling sites (black dots) off New Zealand where shallow-water and deep-water sponges were collected for the incubations. Detailed information about species names and sampling depths are given in Table 1.

**Table 1.** Detailed information about sampling location and species name of all collected sponges. Abbreviation: T = water temperature

| Sample ID | Geographical location | Latitude (°N) | Longitude (°E) | Depth (m) | T (°C) | Species [Class, Family] |
| --- | --- | --- | --- | --- | --- | --- |
| TS002 | Denham Bay, Raoul Island | -29.283 | -177.900 | 6 | 25.0 | *Tedania* sp. indet.<br>[Demospongiae, Tedaniidae] |
| TS004 | Denham Bay, Raoul Island | -29.283 | -177.900 | 6 | 25.0 | *Suberea meandrina* (Kelly, 2015)<br>[Demospongiae, Aplysinellidae] |
| TS005 | Southern Kermadec Ridge (Sponge Ridge) | -35.378 | 178.974 | 1448 | 6.0 | *Farrea raoulensis* (Reiswig & Kelly, 2011)<br>[Hexactinellida, Farreidae] |
| TS006 | Seamount No. 986, off Hawkes Bay Shelf | -39.991 | 178.215 | 828 | 6.0 | *Artemisina* sp. indet.<br>[Demospongiae, Microcionidae] |

### 2.2 *Ex-situ* incubation experiments

For the *ex-situ* incubation experiments aboard R/V *Sonne*, sponges of the same putative species were incubated individually in three closed incubation chambers (Volume: 1.26 L) at *in-situ* temperature in a water bath in a temperature-controlled room (sample ID TS002 and TS004; Table 1, Fig. S1) or in a fridge (sample ID TS005 and TS006; Table 1, Fig. S1). As control, an additional incubation chamber was filled with ambient seawater. Incubations with shallow-water sponges (*Tedania* sp. indet., *Suberea meandrina* (Kelly, 2015); Fig. 2) were conducted as light-incubations, whereas incubations of deep-water sponges (*Farrea raoulensis* (Reiswig & Kelly, 2011), *Artemisina* sp. indet.; Fig. 2) were conducted in the dark.

**Figure 2.**
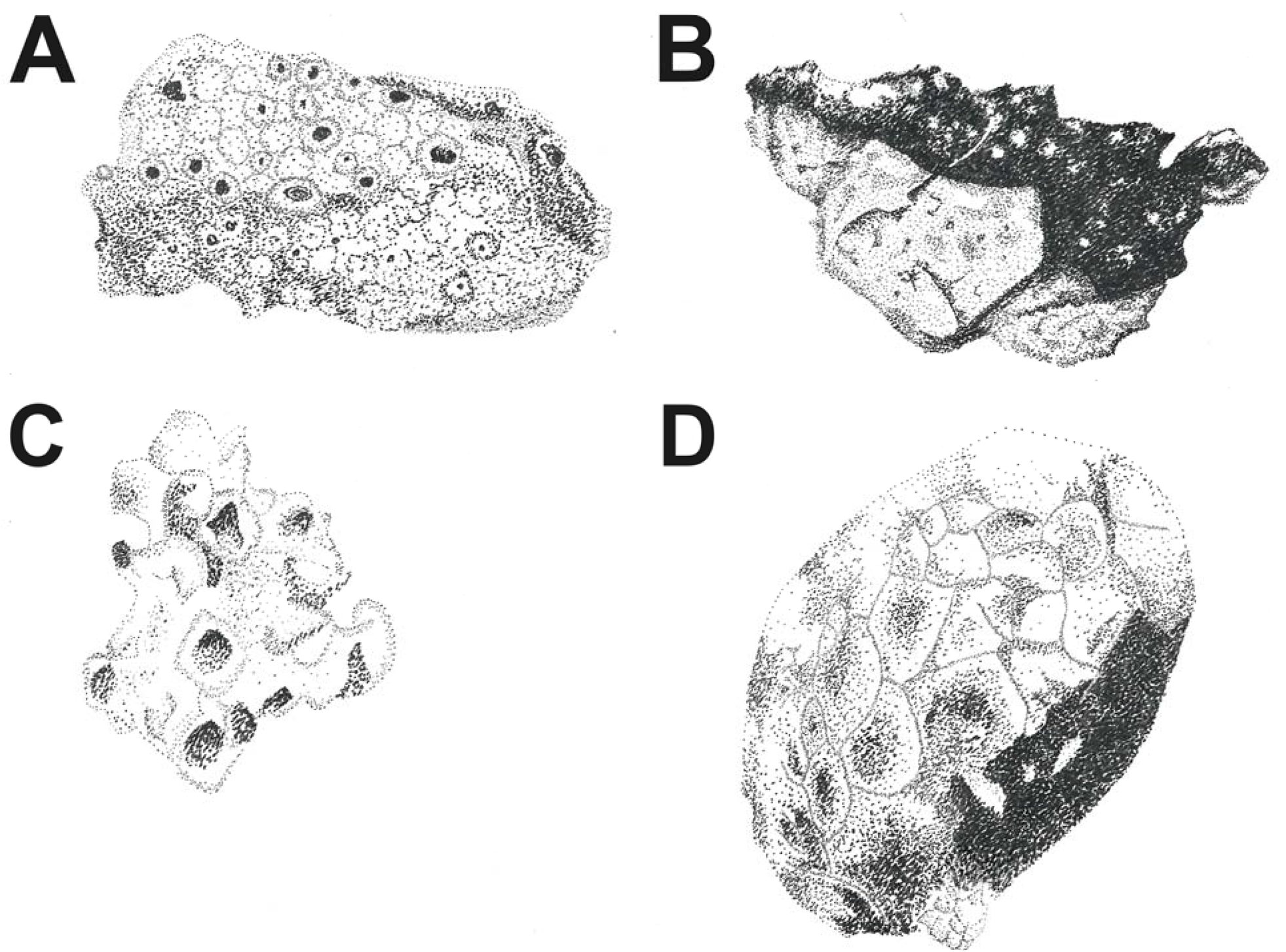
Drawings of the four sponge species A) *Tedania* sp., B) *Suberea meandrina*, C) *Farrea raoulensis*, D) *Artemisina* sp. Sizes of sponge clusters in the drawing are not to scale. Illustrations by Tanja Stratmann.

Every hour for five hours (before the start of the incubation: T0, after 1 h: T1, after 2 h: T2, etc.) triplicate water samples for DOC and TDN were taken, filtered through 0.2 µm filters into acid-washed 30 ml HDPE bottles, acidified to pH 2 with 25% HCl and stored at 4°C until analysis at the University of Oldenburg (Germany). Water samples for colored dissolved organic matter (CDOM) and fluorescent dissolved organic matter (FDOM) analyses were taken at the same time intervals, filtered through 0.2 µm filters into acid-washed, pre-combusted 100 ml amber glass bottles and stored for 1 to 3 d at 4°C until the samples were measured aboard RV *Sonne*.

At the end of the incubations, dimensions (i.e., length *L*, width *W*, and height *H*) of each sponge specimen were measured to the nearest millimeter. Subsamples for microbial community analysis and species identification based on sponge spicules were taken with a scalpel from one sponge per incubation set (Table S1). The subsamples for microbial community analysis were rinsed with sterilized seawater and stored at -80 °C after flash-freezing in liquid nitrogen, whereas subsamples for species identification were kept in ethanol (80%).

### 2.3 Sample processing

#### 2.3.1 Analyses of sponge tissue, fatty acids, and bacterial communities of sponges

Sponge volume (*V*, cm^3^), wet mass (*WM*, g), dry mass (*DM*, g), ash-free dry mass (*AFDM*, g), total carbon (*TC*, % DM)/ δ^13^TC content, total nitrogen (*TN*, % DM)/ δ^15^N content, and organic carbon (org. C, % DM)/ δ^13^org. C content of the sponges were determined as described in detail in (Stratmann et al., 2024). Species-specific conversion factors are presented in Table S2. PLFAs from freeze-dried sponge tissue were extracted following a modified protocol of the Bligh and Dyer extraction method (Bligh and Dyer, 1959; de Kluijver, 2021; de Kluijver et al., 2021) as described in (Stratmann et al., 2024). The bacterial community composition of sponges was extracted and analyzed as described in (Stratmann et al., 2024).

#### 2.3.2 Measurements of DOC, TDN

DOC and TDN concentrations were measured in triplicates by high temperature catalytic combustion with a Shimadzu TOC-VCPH/CPN total organic carbon analyzer coupled to a TNM-1 module. Deep Atlantic Seawater reference material (DSR, D. A. Hansell, University of Miami, USA) was measured during every run to evaluate instrument precision.

#### 2.3.3 Calculation of fluxes and initial net removal rates

Fluxes (*F_i.DOC, TDN_*) of DOC and TDN (µmol g C^-1^ d^-1^) were calculated as:

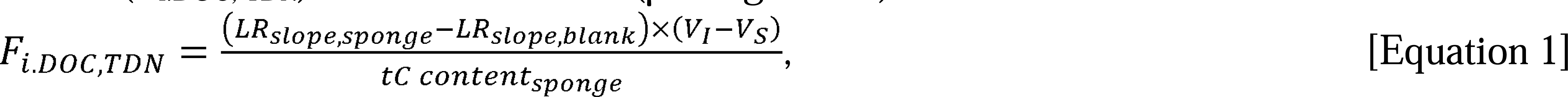

where *LR_slope,sponge_* and *LR_slope,water_* are the estimated slopes of the linear regression analyses done with the concentrations of DOC and TDN (µmol l^-1^) collected over 5 h during the incubations as response (dependent) variable and time as predictor (independent) variable. *tC content_sponge_* corresponds to the org. C content of the incubated sponge (g C), *V_I_* is the volume of the incubation chamber (= 1.26 l) and *V_S_* is the sponge volume (l).

Initial net DOC and TDN removal rates were determined by applying a 2G-model following de Goeij and van Duyl (2007):

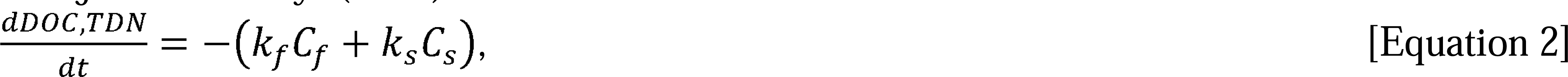

where *C_f_* was the fast removable DOC and TDN fraction, respectively, *C_s_* was the slow removable DOC and TDN fraction, respectively, and *k_f_* and *k_s_* were the corresponding removal constants.

Equation 2 was integrated to present DOC and TDN as a function of time (de Goeij and van Duyl, 2007):

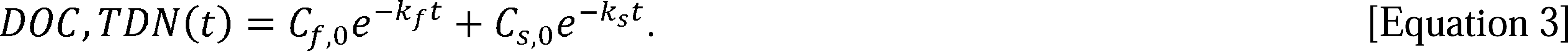

The model variables of Equation 3 were estimated by fitting a non-linear model with the Levenberg-Marquardt algorithm (Moré, 1978) (50 iterations) to the experimental data using the *nlsLM()* function from the *R* package ‘minpack.lm’ (version 1.2-4) (Elzhov et al., 2023) in *R* (version 4.3.0) (R-Core Team, 2022).

Whenever all the model variables could be estimated at a significance level of a = 0.05, the initial DOC and TDN removal rates were calculated following de Goeij and van Duyl (2007) as:

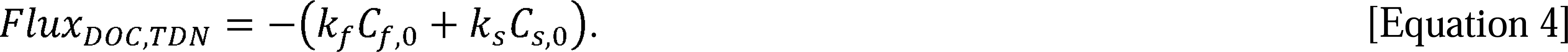

A list with DOC and TDN concentrations at the begin of the *ex-situ* incubations is shown in Table S3.

#### 2.3.4 Measurement of FDOM and parallel factor analysis

Water samples were warmed to room temperature prior to spectrofluorometric measurements in a 1 cm quartz cuvette on an Aqualog spectrofluorometer (HORIBA Scientific, Japan).

Excitation/ emission matrix (EEM) scans were created for excitation wavelengths between 240 and 450 nm with 2 nm increments and for emission wavelengths between 240 and 450 nm with 0.5 nm increments. The excitation and emission bandwidths were 5 nm. EEMs of freshly-produced Milli-Q water served as blank.

EEMs were normalized to Raman Units (RU) after correcting for inner-filter effects (Lakowicz, 2006) and Raman and Rayleigh scatter (Senesi, 1990) using the *DrEEM* toolbox in Matlab version R2017a (MathWorks, Inc.) (Murphy et al., 2013). Subsequently, EEMs were decomposed into independent fluorescent components by parallel factor analysis (PARAFAC; (Bro, 1998; Stedmon et al., 2003)) using the *N-way* toolbox (Andersson and Bro, 2000) in Matlab as described by (Murphy et al., 2013). Spectra of the identified fluorophores were compared with spectra of published FDOM components using the “OpenFluor” (https://openfluor.lablicate.com/) online spectral library for fluorescent, environmental organic compounds (Murphy et al., 2014). Published FDOM spectra were similar to spectra from this study when the Tucker congruence coefficient *r_c_* (Lorenzo-Seva and ten Berge, 2006; Stedmon and Bro, 2008) was >0.95.

### 2.4 Statistical analysis

A significant difference of the average flux of the three incubations of the same sponge species from 0 µmol L^-1^ g C^-1^ d^-1^ was tested by 1-sided Student’s t-test (a = 0.05) in *R* (R-Core Team, 2022) after normality of data was confirmed with a Shapiro-Wilk normality test. When data were not normally distributed, a 1-sample Wilcoxon test (α = 0.05) was performed in *R*.

A change in fluorescence intensity (R.U.) of the identified fluorescing components over time was tested for the three sponge incubations and the control incubation by robust regression analysis (Western, 1995) from the *MASS* package (Venables and Ripley, 2002) in *R*. A change in fluorophore intensity was considered significant for a specific sponge species and fluorophore, when the *F*-statistic (a = 0.05) of the sponge incubation was significant, while the *F*-statistic (a = 0.05) of the control incubation was not significant.

All data are presented as mean ± standard error.

## 3. Results

### 3.1 Composition of sponge tissue

The investigated sponges from New Zealand had a water content of 85.3 ± 1.17% (n = 9) and their dried tissue (n = 8) consisted of 18.3 ± 3.85% TC, 17.5 ± 3.75% org. C, and 4.34 ± 1.02% TN. The lowest TC (3.65 ± 1.62%), org. C (3.03 ± 1.22%), and TN (0.62 ± 0.26%) contents were measured in *F. raoulensis* (n = 3), whereas the sponge species with the highest TC (32.4 ± 1.33%), org. C (31.1 ± 2.01%), and TN (8.28 ± 0.42%) contents was *S. meandrina* (n = 3). The δ^13^org. C-value of the sponges ranged from -17.9‰ (*Artemisina* sp., n = 1) to - 19.7 ± 0.10‰ (*F. raoulensis*, n = 3), the δ^13^TC-value ranged from -18.4‰ (*Artemisina* sp., n = 1) to -20.3 ± 2.55×10^-3^‰ (*F. raoulensis*, n = 3), and the δ^15^N-value ranged from 5.39 ± 0.17‰ (*S. meandrina*, n = 3) to 20.9 ± 2.27‰ (*F. raoulensis*, n = 3) (Fig. 3).

**Figure 3.**
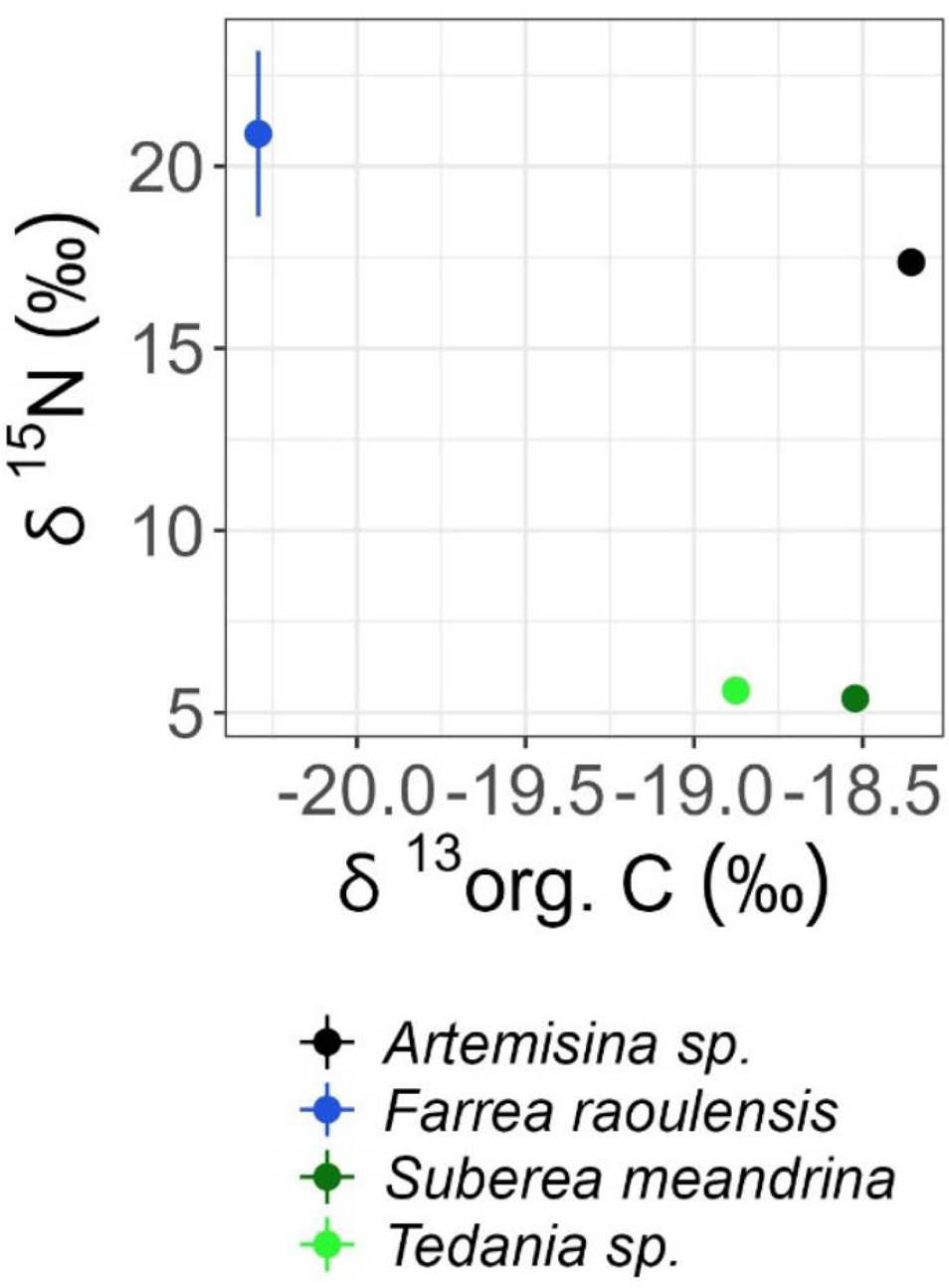
Isotopic composition of carbon (δ^13^org. C, ‰) and nitrogen (δ^15^N, ‰) of sponge species from New Zealand. Error bars indicate 1 standard error.

### 3.2 Dissolved organic matter fluxes

The sponges *Tedania* sp. and *F. raoulensis* released DOC (5.50×10^-2^±9.09×10^-2^ µmol L^-1^ g C^-1^ d^-1^ and 11.3±5.86 µmol L^-1^ g C^-1^ d^-1^, respectively) and TDN (0.26±5.75×10^-2^ µmol L^-1^ g C^-1^ d^-1^ and 31.5±8.22 µmol L^-1^ g C^-1^ d^-1^, respectively), though only the TDN flux of *Tedania* sp. was significantly different from 0 (Table 2). *S. meandrina* took up DOC (−0.12± 5.16×10^-2^ µmol L^-1^ g C^-1^ d^-1^) and TDN (−2.49±0.94 µmol L^-1^ g C^-1^ d^-1^), again not at significant rates (Table 2).

**Table 2.** Results of 1-sided Student’s t-tests (α = 0.05) and 1-sample Wilcoxon tests (α = 0.05) to assess whether DOC and TDN fluxes (µmol g org. C^-1^ d^-1^) were significantly different from 0 (H_0_: µ = 0, H_1_: µ -;;/ 0). Symbols: \**p*-value≤0.05, L*p*-value≤0.01

| Sample ID | Flux type | t-value | df | V | p-value | Mean | 95% Confidence interval |
| --- | --- | --- | --- | --- | --- | --- | --- |
| *Tedania sp.* ($n = 3$) | DOC | 0.61 | 2 | | 0.61 | 0.056 | -0.34, 0.45 |
|  | TDN | 4.48 | 2 |  | 0.05* | 0.26 | 0.01, 0.50 |
| *Suberea meandrina* ($n = 3$) | DOC | -2.35 | 2 | | 0.14 | -0.12 | -0.34, 0.10 |
|  | TDN |  |  | 0 | 0.25 |  |  |
| *Farrea raoulensis* ($n = 3$) | DOC | 1.93 | 2 | | 0.19 | 11.3 | -13.89, 36.5 |
|  | TDN | 3.83 | 2 |  | 0.06 | 31.5 | -3.90, 66.9 |

No initial net DOC and TDN removal rates could be calculated, because the estimated parameters were not significant for any incubation (Table S4).

### 3.3 Fluorescent dissolved organic matter composition

The PARAFAC model could be validated for four different fluorophores (Fig. 4). Fluorophores C1 (λ_Ex_= 294, λ_Em_ = 350) and C2 (λ_Ex_ = 280, λ_Em_ = 337) were similar to peak T and tryptophan- and protein-like (Coble, 2007, 1996). Fluorophore C3 (λ_Ex_ = <260/ 332, λ_Em_ = 446) was similar to peaks A and C and humic-like (Coble, 2007, 1996), whereas fluorophore C4 (λ_Ex_ = 274, λ_Em_ = <300) was similar to peak B and tyrosine- and protein-like (Coble, 2007, 1996).

**Figure 4.**
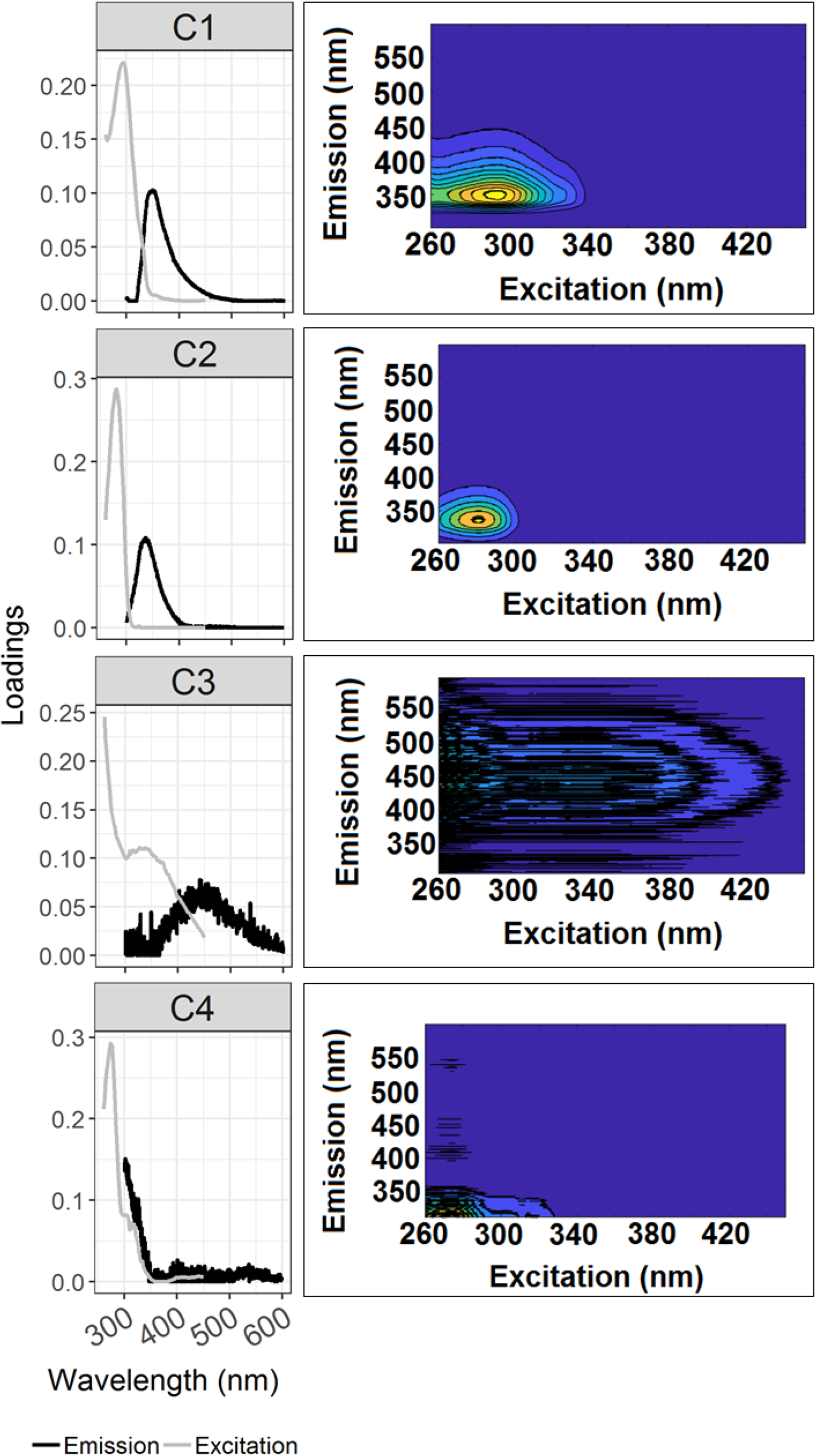
Spectral loadings (Excitation/ Emission; left panels) of the fluorophores C1, C2, C3, and C4 that were identified by PARAFAC of all samples and their corresponding emission excitation scans (right panels).

A comparison of FDOM spectra of the four fluorophores with published FDOM spectra from the “OpenFluor” online spectral library resulted in 201 comparable components (C1: 39, C2: 62, C3: 52, C4: 48) at *r_c_*≥0.95 (Table S5).

The fluorescence intensity (R.U.) of fluorophore C1 decreased significantly over time in incubations with *S. meandrina* (F_1,16_ = 5.77, p = 0.03) (Fig. 5, Table S6). At the same time, the fluorescence intensity of fluorophores C2 and C4 increased over time [F_1,16_ (C2) = 5.60, p = 0.03; F_1,16_ (C4) = 9.17, p = 0.008). In comparison, the fluorescence intensity of C3 in incubations of *Tedania* sp. increased significantly over time (F_1,16_ = 19.2, p = 0.0005), whereas the fluorescence intensity of C4 in the same incubations decreased (F_1,16_ = 4.27, p = 0.05).

**Figure 5.**
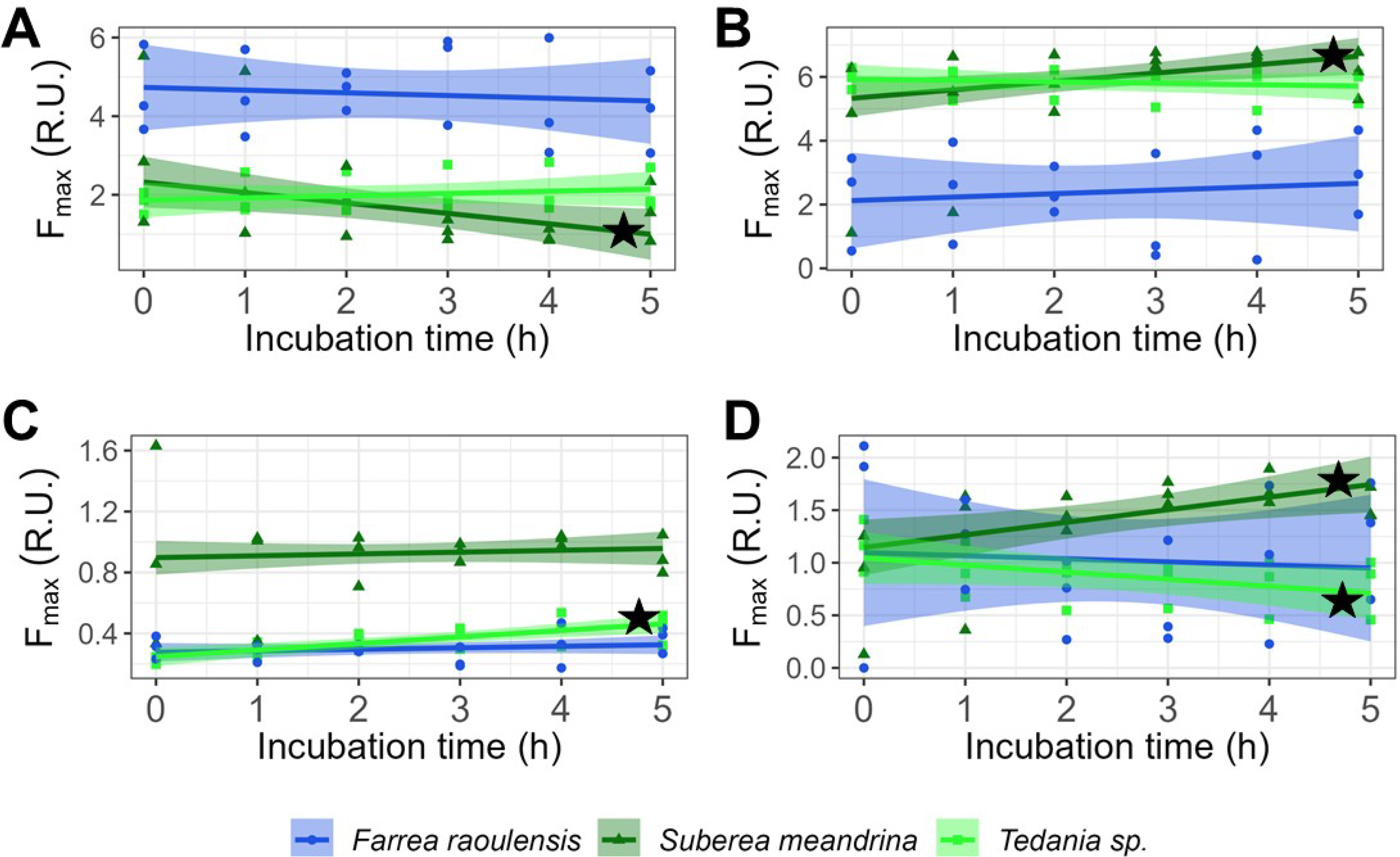
Relationship of maximum fluorescence intensity F_max_ (Raman Units R.U.) to incubation time (h) for the fluorophores (A) C1, (B) C2, (C) C3, and (D) C4 that were identified in the PARAFAC model (Fig. 4).

Black asterisks mark robust regression analysis results that were significant at a significance level a = 0.05 (Table S6).

### 3.4 Fatty acid composition of sponges

The sponges showed species-dependent differences in their PLFA compositions (Fig. 6; Table S3). Between 12.5% (*S. meandrina*) and 46.8% (*F. raoulensis*) of the PLFAs found in sponges consisted of long-chain fatty acids (LCFA, i.e., fatty acid with ≥24 C atoms). The other PLFA classes present in the sponges were branched fatty acids (4.48 ± 4.48%), cyclic fatty acids (1.80 ± 1.12%), highly unsaturated fatty acids (HUFA, i.e., fatty acids with ≥4 double bonds; 3.50 ± 2.89%), methyl-fatty acids (5.44 ± 2.24%), monosaturated fatty acids (MUFA; 18.7 ± 6.24%), polyunsaturated fatty acids (PUFAs, i.e., fatty acids with ≥2 double bonds; 0.49 ± 0.31%), saturated fatty acids (SFA; 22.9 ± 2.72%), and undefined fatty acids (10.3 ± 5.66%).

**Figure 6.**
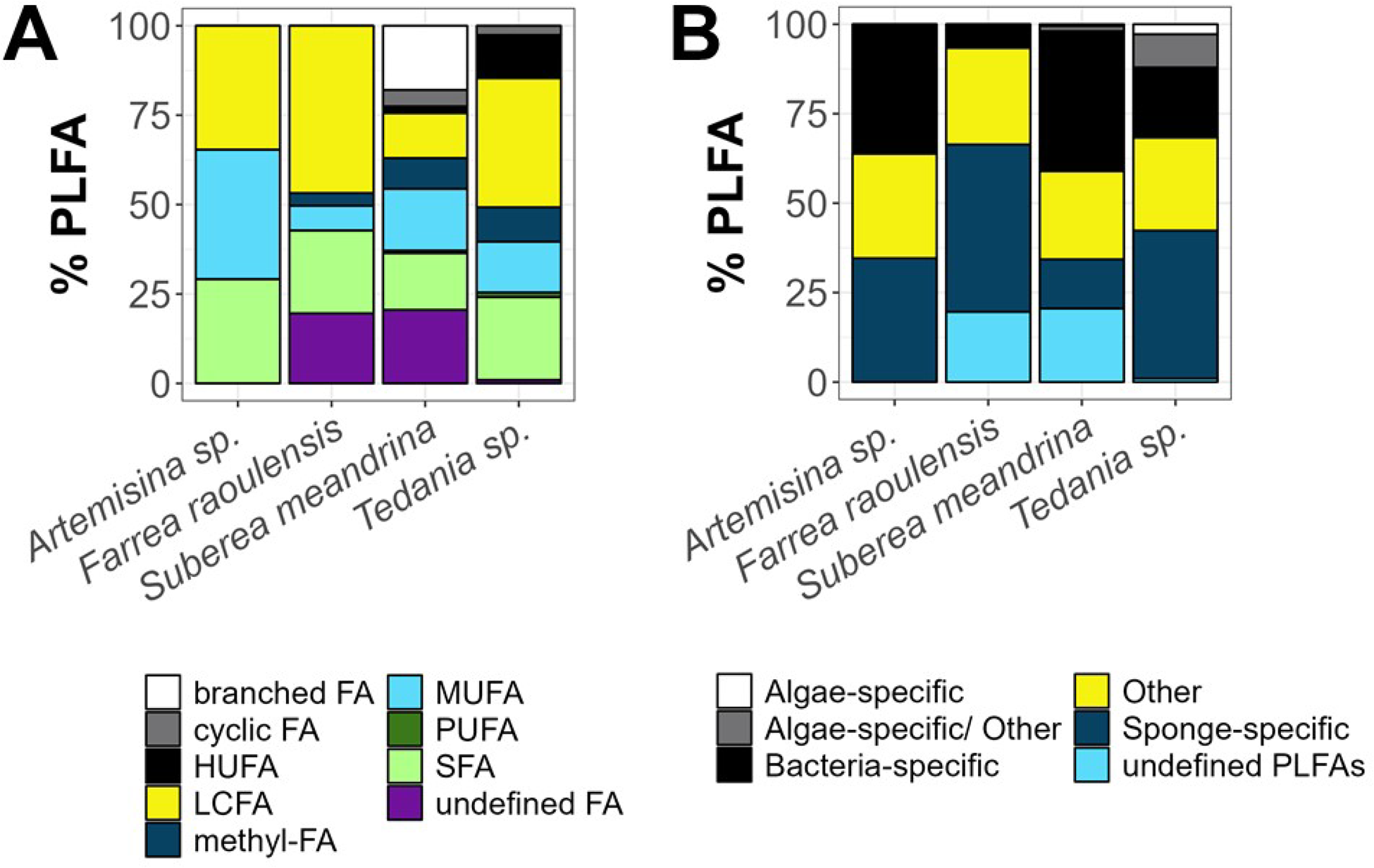
(**A**) Contribution (%) of individual phospholipid-derived fatty acid (PLFA) classes to the total concentrations in sponges. (**B**) Contribution (%) of individual PLFA categories to total PLFA concentrations in sponges. Abbreviations: branched FA = branched fatty acid, cyclic FA = cyclic fatty acid, HUFA = highly unsaturated fatty acid, LCFA = long-chain fatty acid, methyl-FA = methyl-fatty acid, MUFA = monounsaturated fatty acid, PUFA = polyunsaturated fatty acid, SFA = saturated fatty acid.

Most isolated PLFAs were sponge-specific PLFAs (min: 13.7% of total PLFA concentration, *S. meandrina*; max: 46.8%, *F. raoulensis*) (Fig. 6B), followed by bacteria-specific PLFAs (min: 6.61%, *F. raoulensis*; max: 39.1%, *S. meandrina*), and others (min: 24.7%, *S. meandrina*; max: 29.2%, *Artemisina* sp.). Between 0.00% (*Artemisina* sp.) and 20.6% (*S. meandrina*) of the PLFAs were unidentified.

### 3.5 Composition of sponge-associated bacterial community

The bacterial communities of *Artemisina* sp., *S. meandrina*, and *Tedania* sp. were dominated by Proteobacteria (min: 21.7% ASVs *S. meandrina*; max: 54.0% ASVs *Tedania* sp.) and Chloroflexota (min: 30.1% ASVs *S. meandrina*; max: 51.2% ASVs *Artemisina* sp.) (Fig. 7, Table S6). The bacterial community of *F. raoulensis* was not enriched in Chloroflexota to a comparable extent, but consisted to 97.6% of Proteobacteria. The third and fourth most abundant bacterial phyla in *S. meandrina* were Actinobacteria (14.9% ASVs) and Acidobacteria (12.1% ASVs) (Fig. 7).

**Figure 7.**
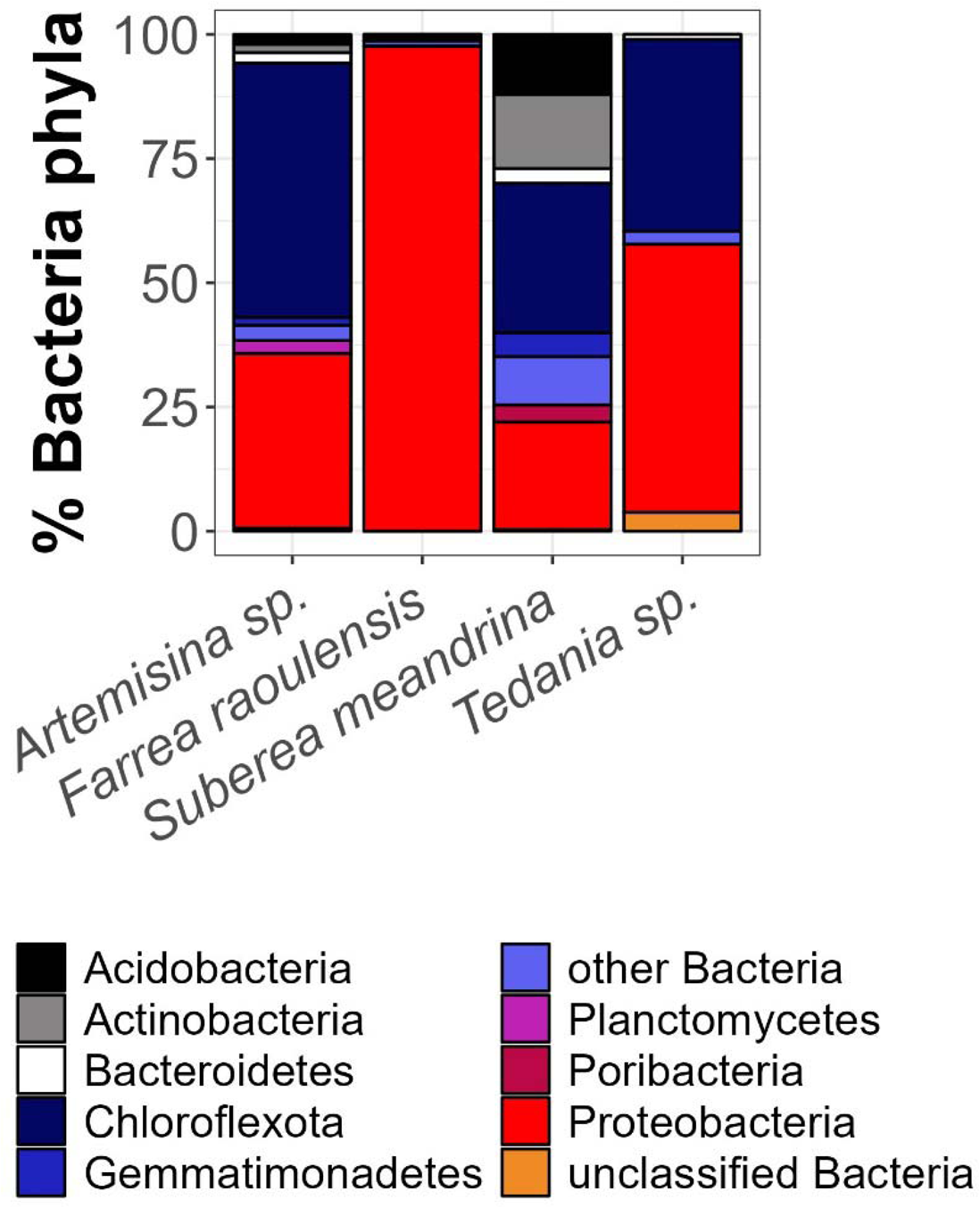
Relative abundance of bacteria phyla (%) in the four sponges. The eight most abundant bacteria phyla are indicated by names, others are aggregated under the category “other Bacteria” and “unclassified Bacteria”.

## 4. Discussion

### 4.1 Dissolved organic matter fluxes

Over the last 20 years, numerous studies have shown that sponges are capable of taking up DOM (e.g., Bart et al., 2020; de Goeij et al., 2013, 2008b, 2008a; Morganti et al., 2017; Ribes et al., 2023; Rix et al., 2017; Yahel et al., 2003). In fact, it has been proposed that sponges use DOM as their main energy source for basal metabolic processes, while they use bacterioplankton and other particulate organic matter (POM) for anabolic processes, such as somatic growth, cell turnover, and reproduction (Bart et al., 2020). Unfortunately, we did not measure the concentrations of bacterioplankton or POM in the incubations and therefore could not calculate any uptake rates, but estimations of DOC and TDN fluxes showed no significant assimilation of DOC, and even a significant release of TDN by *Tedania* sp.

Some sponge species require a minimum DOC concentration in the ambient seawater (e.g., 79 µmol C l^-1^ seawater for *Xestospongia muta* (Schmidt, 1870) (Wooster et al., 2019); 74/80/198 µmol C l^-1^ seawater for *Agelas oroides* (Schmidt, 1864), *Crambe crambe* (Schmidt, 1862), *Chondrosia reniformis* Nardo, 1847, *Dysidea avara* (Schmidt, 1862) (Morganti et al., 2017; Ribes et al., 2023); 80 µmol C l^-1^ for *Petrosia* (*Petrosia*) *ficiformis* (Poiret, 1789)

(Morganti et al., 2017)) to be able to assimilate DOM. In the incubation settings here, the initial DOC concentrations ranged from 141 to 297 µmol C l^-1^ seawater (Table S3) and therefore surpassed most of the DOC thresholds. Hence, if the sponges were capable of exploiting the DOC pool, they should have taken it up. However, the lack of significant DOC assimilation by sponges in this study compared to sponges in other studies might also be related to the inconsistent operational definition of DOC (<0.2 μm pore size vs. <0.7 μm pore size). Several studies that measured DOC consumption used a pore size of <0.7 μm (e.g., (Achlatis et al., 2019; Bart et al., 2021; de Goeij et al., 2013; Hoer et al., 2018; Morganti et al., 2017)), which includes colloids and particles, whereas we used a pore size of <0.2 μm which should only allow truly dissolved DOC to pass (Verdugo et al., 2004). Hence, the sponges in this study are either unable to take up DOC or they incorporate colloidal DOC, but not truly dissolved DOC.

### 4.2 Changes in fluorescent dissolved organic matter composition

The tryptophan-like fluorophores identified by the PARAFAC model can be of autochthonous origin (Kellerman et al., 2015) and microbial-derived (Walker et al., 2013), may have a low molecular weight (Wagner et al., 2015), and be part of the labile or semi-labile fraction of DOM (Davis and Benner, 2007). Humic-like fluorophores may originate from humic acids that were released during microbial processing of DOM (Stedmon and Markager, 2005; Yamashita et al., 2010) or may be generated by photo-production (Chen et al., 2010; Miranda et al., 2020).

Tyrosine-like fluorophores originate from aromatic amino acids (Yamashita and Tanoue, 2003) and are of labile or semi-labile nature (Davis and Benner, 2007) like the tryptophane-like fluorophores. These fluorophores are often degradation products (Romero et al., 2017) of small peptides or proteins (Yamashita and Tanoue, 2003).

In the incubations, *S. meandrina* degraded the tryptophan-like fluorophore C1 and/ or transformed it into a fluorophore C2 with different excitation and emission maxima that still resembled a protein- or tryptophan-like fluorescence pattern, while it also produced a tyrosine-like fluorophore C4. In comparison, *Tedania* sp. degraded the tyrosine-like fluorophore C4 and excreted a humic-like fluorophore C3.

The removal of protein-like fluorophores and release of humic-like fluorophores was reported for the Mediterranean sponges *C. reniformis*, *A. oroides*, *C. crambe*, and *D. avara*, that occasionally also excreted tryptophan-like fluorophores (Ribes et al., 2023). In fact, an untargeted metabolic analysis of exhalent and inhalant seawater samples of the sponges *Ircinia campana* (Lamarck, 1814) and *Spheciospongia vesparium* (Lamarck, 1815) from the Florida Keys detected an increase in the concentration of tryptophan and tyrosine in exhalent compared to inhalant seawater besides the release of 241 other metabolites (Fiore et al., 2017). The authors even estimated that each sponge would excrete up to 1 nmol tryptophane l^-1^ seawater and 1 nmol phenylalanine l^-1^ seawater. Studies using ultrahigh-resolution mass spectrometry furthermore revealed that the sponges *Melophlus sarasinorum* Thiele, 1899, *Rhabdastrella globostellata* (Carter, 1883), and *S. vesparium* preferentially removed low-molecular mass DOM compounds (Hildebrand et al., 2022; Letourneau et al., 2020). The chemical composition of these removed compounds, however, differed between sponge species. *Melophlus sarasinorum* and *R. globostellata* targeted compounds with high C/N and high H/C ratios (Hildebrand et al., 2022), whereas *S. vesparium* depleted mainly compounds with a high N/C ratio and a low number of oxygen atoms (Letourneau et al., 2020). Hence, the shallow-water sponges in this study (*Tedania* sp., *S. meandrina*) alter the DOM composition of the seawater in a sponge-species-specific way and do not produce a “characteristic ‘sponge holobiont signal’” as proposed by Fiore et al., (2017).

### 4.3 Microbial community of the sponge holobiont

Based on the microbial community composition of the sponges, *F. raoulensis* is a LMA sponge with a dominance of Proteobacteria, whereas the high abundance of Acidobacteria, Actinobacteria and Chloroflexota in *S. meandrina* suggests this species may be a HMA sponge (Busch et al., 2022). Some species belonging to the genus *Tedania* have been predicted to be LMA sponges (Busch et al., 2022; Moitinho-Silva et al., 2017), but the *Tedania* sp. individual shown here displays more of a HMA profile, with an enrichment in Chloroflexota. The *Artemisina* sp. individual also displays a HMA profile with comparably high relative abundances of Chloroflexota.

Bacteria of the phyla Bacteroidetes (Grondin et al., 2017; O’Brien et al., 2023), Poribacteria (Jahn et al., 2016; Kamke et al., 2013; O’Brien et al., 2023), and Chloroflexota (Bayer et al., 2018; Colatriano et al., 2018; Landry et al., 2017) are able to degrade DOM. Indeed, as symbionts of the sponge *Plakortis angulospiculatus* (Carter, 1879), members of the phyla Poribacteria and Chloroflexiota, together with members of the phyla PAUC34F, Proteobacteria, and Nitrospirota, were active consumers of DOM in a DNA-stable isotope probing experiment (Campana et al., 2021). The authors proposed that a PAUC34f bacterium and bacteria of the genus *Endozoicomonas* (phylum Proteobacteria) were the first ones to consume and incorporate DOM, followed by a Poribacterium, a Nitrospira bacterium and three Chloroflexota bacteria.

Hence, we speculate that in the incubation experiments with New Zealand sponges, Chloroflexota and Poribacteria may have been involved in degradation and transformation of DOM, too. Both phyla were present in *S. meandrina* and Chloroflexota was additionally detected in *Tedania* sp. (Table S8). The contribution of Chloroflexota to the overall sponge-associated bacteria in *F. raoulensis*, in comparison, was <1% which might give one reason why we did not observe a significant change in DOC/ TDN concentration or FDOM composition in incubations with this sponge species.

### 4.4 Fatty acid composition of deep-sea sponges

The fatty acid composition of the four analyzed sponges differed quite substantially. The deep-sea hexactinellid *F. raoulensis* contained seven different identified PLFAs and the shallow-water demosponges *Tedania* sp. and *S. meandrina* had a diverse range of 22 and 29 identified PLFAs, respectively. In comparison, the PFLA pool of the deep-sea demosponge *Artemisina* sp. consisted of only three different PLFAs (Table S7).

The class of Hexactinellida sponges is characterized by a reduced diversity of PLFAs (Stratmann et al., 2024; Thiel et al., 2002a), very long C chains of 28 to 32 C atoms, and a lack of C chains with 24 to 28 C atoms (Thiel et al., 2002a). A comparison of the PLFA composition of *F. raoulensis* with the composition of two specimens of the Sceptrulophora cluster from Stratmann et al., (2024) showed a very close resemblance in the relative abundance of the individual PLFAs, but *F. raoulensis*’ PLFA composition deviated from the PLFA composition of *Saccocalyx tetractinus* (Reiswig and Kelly, 2018). The latter contained almost 40% C29:2ω?, but <10% C30:3ω? (Stratmann et al., 2024), whereas *F. raoulensis* had <10% C29:2ω? and about 25% C30:3ω?. Hence, hexactinellid sponges of the same genera that originate from the same region seem to have a very similar PLFA composition in contrast to demosponges. de Kluijver et al., (2021) investigated the fatty acid profiles of several boreal and Arctic demosponges of the genera *Geodia* and *Stelletta* and found that *G. barretti*, *Geodia atlantica* (Stephens, 1915), and *Geodia hentscheli* Cárdenas, Rapp, Schander & Tendal, 2010 synthesize similar LCFA in different relative abundances, whereas *Geodia parva* Hansen, 1885 and *Stelletta rhaphidiophora* Hentschel, 1929 synthesize distinct LCFA.

These species-specific differences in fatty acid composition in Demospongiae instead of genus-specific differences like in Hexactinellida hampers direct testing whether temperature affects fatty acid composition of sponges. Nevertheless, we compared the PLFA composition of two species of the genus *Tedania* that were collected during the same period in waters around New Zealand: *Tedania* sp. from Raoul Island (6 m water depth; this paper) lived at an ambient temperature of 25°C, while *Tedania* sp. from Pegasus canyon, off Canterbury (852 m depth; (Stratmann et al., 2024)) lived at 6°C. Following observations by Bennett et al., (2018), sponges may adapt to increasing water temperatures by increasing the mean chain length of their fatty acids and increasing the degree of unsaturation of fatty acids when they are phototrophic sponges or decreasing the degree of unsaturation of fatty acids when they are heterotrophic sponges.

These adaptations could not be found in *Tedania* sp. The species living at the lower temperature had less short-chain PLFAs than the species living at higher temperature and only 0.5% more unsaturated PLFA. Hence, either the temperature adaptation in *Tedania* sp. is less pronounced than in *Phyllospongia foliascens* (Pallas, 1766), *Cymbastela coralliophila* Hooper & Bergquist, 1992, *R. odorabile*, and *Stylissa flabelliformis* (Hentschel, 1912), or the differences in fatty acid composition among species is stronger and cover any environmental effects.

## 5. Conclusions

In this study, we investigated the role of shallow- and deep-water sponges originating from New Zealand in the processing of DOM. *Suberea meandrina* takes up insignificant amounts of DOC and TDN, whereupon bacterial members of its holobiont (likely from the phyla Chloroflexota and Poribacteria) may be involved in the degradation of tryptophan-like fluorophores to a tryptophan-like fluorophore with a different chemical composition. At the same time, the sponge holobiont produces tyrosine-like fluorophores. *Tedania* sp. releases significant amounts of TDN, likely due to the production of humic-like fluorophores, while the holobiont, particularly its bacteria member Chloroflexota, may contribute to the degradation of tyrosine-like fluorophores. Furthermore, the PLFA composition of different Tedania species maybe less adapted to temperature and more dependent on species-specific differences or other environmental factors. *Farrea raoulensis* is not involved in DOM processing which could be related to a lack of Chloroflexota. They have a reduced diversity of PLFAs which is very similar to the PLFA composition of other members of the *Farrea* genus, but contain numerous different identified fatty acids than *Artemisina* sp.

## Funding

This research received funding from the Royal Netherlands Academy of Arts and Sciences (KNAW, The Netherlands) to TS (Academy Ecology Grant 2017) and from the Dutch Research Council (NWO, The Netherlands) to TS (NWO-Rubicon grant no. 019.182EN.012, NWO-Talent program Veni grant no. VI.Veni.212.211). AdK was supported by the SpongGES project of the European Union’s Horizon 2020 research and innovation program (grant no. 679849). PS received funding by the Federal Ministry of Education and Research (BMBF) for the cruise SO254, grant no. 03G0254A, PORIBACNEWZ.

Specimens were collected as part of the project “PoriBacNewZ” by the Institut für Chemie und Biologie des Meeres (ICBM), University Oldenburg on the German flagship RV *Sonne*, using the ROV Kiel 6000 (GEOMAR) with participation and funding from GEOMAR, DSMZ, LMU, NIOZ, NIWA, and ETH-Zurich. NIWA voyage participation was funded through the MBIE SSIF Enhancing Collections project.

## CRediT Roles

**Tanja Stratmann**: Conceptualization, Data curation, Formal Analysis, Funding acquisition, Investigation, Methodology, Project administration, Resources, Software, Validation, Visualization, Writing – original draft; **Mario Miranda**: Formal Analysis, Investigation, Methodology, Writing – review & editing; **Anna de Kluijver**: Investigation, Methodology; **Kathrin Busch**: Data curation, Investigation, Methodology, Writing – review & editing; **Michelle Kelly**: Taxonomic identification, Writing – review & editing; **Sadie Mills**: Data curation, Resources, Validation, Writing – review & editing; **Peter Schupp**: Funding acquisition, Resources, Supervision, Writing – review & editing.

## Declaration of competing interests

There are no conflicts of interest declared for this submission.

## Data availability

All data presented in this manuscript are available at Pangaea (**XXX**) and on NCBI (accession numbers: PRJNA1102912; http://www.ncbi.nlm.nih.gov/bioproject/1102912).

Sample collection was carried out under the “Application for consent to conduct marine scientific research in areas under national jurisdiction of New Zealand (dated 7.6.2016).” Sponge collection at Raoul Island was performed under the “Authorization to undertake specified scientific study within a marine reserve” (authorization number: 53535-MAR, dated 22.12.2016).

## Acknowledgements

The authors thank chief scientist Prof. Meinhard Simon (University of Oldenburg), the captain and crew of RV *Sonne*, and the ROV Kiel 6000 team from Geomar (Kiel) for their excellent support during research cruise SO254. The authors acknowledge the analytical support from Peter van Breugel (NIOZ) and from Klaas Nierop (Utrecht University). They further acknowledge Thorsten Dittmar, Ina Ulber, and Matthias Friebe (University of Oldenburg) for conducting the TDN/ DOC analyses.

## Notes

### Competing Interest Statement

The authors have declared no competing interest.

